# Black soldier fly (*Hermetia Illucens*) larvae meal fermented with *Lactobacillus bulgaricus* as probiotic feed

**DOI:** 10.64898/2025.12.15.694476

**Authors:** Piero-Alexander Medina-Herrera, Anggeli-Solansh Huacasi-Bellido, Mariajosé Roldán-Apaza, Edith-Evelin Mendoza-Mamani, Kevin Tejada-Meza

**Affiliations:** Catholic University of Santa María

**Keywords:** Black soldier fly larvae meal, BSFLM, Probiotic meal, Soy flour, Fermentation, Probiotic, Antimicrobial, Lactobacillus, Sustainable livestock farming, Circular economy

## Abstract

The overuse of antibiotics in livestock is leading to the development of super-resistant pathogens, necessitating the production of probiotic feed for livestock that is also sustainable. Therefore, the study aims to ferment five mixtures of black soldier fly larvae meal (BSFLM) and soy in different proportions with the probiotic *Lactobacillus bulgaricus* in order to determine which allows the greatest growth of this probiotic bacteria and represent a new feed alternative for sustainable livestock farming. To achieve this, the growth curve of *Lactobacillus bulgaricus* in MRS broth was determined. In the exponential growth phase, this broth was added to the five formulations of BSFLM and soy, and finally, the CFU produced by each formulation was determined. It was found that the concentration of BSFLM has a statistically significant effect (p < 0.05) on the concentration of *Lactobacillus bulgaricus*, with twice as many CFU of the probiotic obtained in the pure HLMSN treatment than in the pure soy treatment. These results reveal that *Lactobacillus bulgaricus* is capable of metabolizing and producing greater biomass with BSFLM as a substrate instead of soy, so this formulation can be used in subsequent studies for *in vitro* and *in vivo* antimicrobial testing.

## INTRODUCTION

Antimicrobial resistance (AMR) is one of the major global challenges of the 21st century. The increasing prevalence of multidrug-resistant pathogens represents a worldwide public health problem, primarily driven by the excessive and inappropriate use of antibiotics (Bhardwaj et al., 2022; Akram et al., 2023), which promotes the development of resistance mechanisms and progressively reduces the effectiveness of available antimicrobial therapies (Akram et al., 2023). Antibiotics have been extensively used in both human and animal populations; in livestock, they are commonly applied not only to treat bacterial infections but also as growth promoters (Jian et al., 2021). Consequently, pathogens acquire and disseminate antimicrobial resistance genes (ARGs), which may persist in the environment even after microbial death, and spread through horizontal and vertical gene transfer mechanisms, thereby exacerbating the current global health crisis (Ajayi et al., 2024).

In Peru, several studies have reported a significant increase in multidrug-resistant pathogens in animal populations (Cabrera et al., 2023). The presence of resistant pathogens within livestock microbiota raises concerns regarding their transmission through animal-derived food products (Hassani et al., 2022; Negi and Sharma, 2024), as confirmed by microbiological analyses of artisanal cheeses produced in Canta, Lima, Peru (Reyes et al., 2023). These findings highlight the close relationship between antibiotic use in livestock production and its indirect adverse effects on human health, including increased morbidity, mortality, disease duration, comorbidity development, and the emergence of epidemic outbreaks (Bava et al., 2024).

In addition to public health concerns, current livestock meat production systems have been recognized as environmentally unsustainable and highly damaging to ecosystems (Kumar et al., 2022), which has stimulated growing interest in the development of alternative and sustainable food sources (Xiong, 2023; Caputo et al., 2024). In this context, insect-based feeds have emerged as a promising alternative for livestock nutrition (Sogari et al., 2019; Sogari et al., 2023). Insects are characterized by a low environmental footprint, high growth rates, simple rearing requirements, and an exceptional capacity to convert a wide range of organic waste into high-value, nutrient-dense products, thereby contributing to circular economy models (Elahi et al., 2022). Moreover, insects provide substantial nutritional benefits for livestock, exhibiting high levels of protein, lipids, essential amino acids, and minerals (Fu et al., 2024). Depending on rearing conditions, protein content in insects such as the black soldier fly (*Hermetia illucens*) larvae may range from 35–60%, with lipid contents between 25–50%, along with a complete profile of essential amino acids and minerals (Smets et al., 2020; Lu et al., 2022).

Beyond their nutritional value, insect-enriched feeds have been reported to exert positive effects on animal health. Structural components such as chitin have been shown to stimulate the immune system and exhibit antimicrobial activity (Rahman et al., 2023; Manniello et al., 2021). Furthermore, insect-based substrates can be subjected to probiotic fermentation processes, enhancing their antimicrobial properties against intestinal pathogens such as *Salmonella, Klebsiella*, and *Escherichia coli* (Islam and Yang, 2017). Studies evaluating antimicrobial activity consistently emphasize the importance of conducting antimicrobial assays using insect-based fermented feeds inoculated with probiotic microorganisms, particularly *Lactobacillus* spp. (Sarica et al., 2024; Song et al., 2015).

Based on the aforementioned evidence, black soldier fly larvae meal (BSFLM) is hypothesized to be a suitable substrate for probiotic growth. Accordingly, the present study aimed to evaluate the ability of *Lactobacillus bulgaricus*, a well-established gastrointestinal probiotic, to utilize BSFLM as an optimal substrate in comparison with formulations containing varying concentrations of soybean meal. Soybean meal is widely used in animal feed worldwide; however, its large-scale cultivation has become increasingly unsustainable (Pendrill et al., 2022). To achieve this objective, the microbial growth kinetics of *Lactobacillus bulgaricus* were first assessed in de Man, Rogosa and Sharpe (MRS) broth. Subsequently, five different formulations with varying proportions of BSFLM and soybean meal were prepared and inoculated. After incubation, aliquots were collected, serially diluted, and plated on MRS agar. Colony-forming units (CFU) were quantified for each formulation, and the resulting data were analyzed using a completely randomized design (CRD) to determine whether BSFLM concentration exerted a significant effect on CFU per gram.

## MATERIALS AND METHODS

### Geographical location

All experimental procedures described below were conducted between January and June 2024 in the multidisciplinary research laboratories of the Universidad Católica de Santa María (Arequipa, Arequipa, Peru), located at 2,335 m above sea level, with an average ambient temperature of 22 °C.

### Acquisition of black soldier fly larvae meal (BSFLM) and soybean meal

Black soldier fly larvae meal (BSFLM) was acquired from KAWAT S.A.C., located in Rioja, Nueva Cajamarca, Peru. Soybean meal was obtained from a local supplier in Arequipa, Arequipa, Peru.

### Source and preservation of *Lactobacillus bulgaricus*

The bacterial strain was provided by the Microbiology Laboratory of the Universidad Católica de Santa María (Arequipa, Arequipa, Peru). The stock culture of *Lactobacillus bulgaricus*, a widely studied lactic acid bacterium (LAB), was prepared on de Man, Rogosa and Sharpe (MRS) agar plates (HiMedia, India) and incubated at 37 °C for 24 h. After incubation, the plates were stored at 4 °C until further use.

### Determination of the growth curve by visible spectrophotometry

An initial inoculum was prepared by culturing *Lactobacillus bulgaricus* in 5 mL of MRS broth (HiMedia) to achieve a high biomass concentration suitable for subsequent growth curve analysis. The culture was incubated overnight at 33 °C.

On the following day, 100 mL of fresh MRS broth (HiMedia) was prepared for growth curve determination. Prior to inoculation, a blank solution was established by measuring the absorbance of 5 mL of uninoculated MRS broth using visible-range spectrophotometry at 480 nm. Subsequently, 5 mL of the overnight-incubated inoculum was added to the broth, vigorously mixed, and incubated at 37 °C.

During a 20 h incubation period, 5 mL aliquots were withdrawn every 2 h to measure absorbance and construct the microbial growth curve. These aliquots were also used for viable cell determination by the improved Neubauer chamber method, as described below.

### Viable cell counting using the improved Neubauer chamber method

Simultaneously with absorbance measurements, the 5 mL aliquots collected from the culture were refrigerated at 4 °C and subsequently used to quantify viable cells using the improved Neubauer chamber method (Zhang et al., 2020). This method is referred to as “improved” because it yields results that more closely approximate those obtained using specialized automated cell-counting instruments.

As in the conventional method, dead cells are stained blue, whereas viable cells remain relatively transparent. However, in the optimized method, cells located along all four chamber borders are counted rather than only the upper and left borders. The total border count is then divided by two, and subsequent calculations are performed according to the applied dilution factor.

### Preparation of BSFLM and soybean meal formulations for fermentation

Once the time point corresponding to maximum *Lactobacillus bulgaricus* growth in MRS broth under the previously described conditions was identified, five different fermentation formulations were prepared according to the proportions detailed in Table 1. Each formulation had a total mass of 28 g, of which 10 g corresponded to MRS broth and the remaining fraction consisted exclusively of solid substrates (BSFLM and soybean meal).

**Table 1.** Composition of each formulation of HLMSN and soybean meal.

| Trial | % of soybean meal | % of BSFLM |
| --- | --- | --- |
| 1 | 0 | 100 |
| 2 | 25 | 75 |
| 3 | 50 | 50 |
| 4 | 75 | 25 |
| 5 | 100 | 0 |

After weighing and homogenizing both flours, MRS broth containing *Lactobacillus bulgaricus* in the exponential growth phase was added. Each system was then allowed to ferment for 48 h at 33 °C under closed-system conditions.

### Serial dilutions and plate counting of colony-forming units (CFU)

After 48 h of fermentation, each closed fermenter was opened and serial dilutions were prepared using 1% saline solution in Falcon tubes. Dilution factors of 10^−1^, 10^−2^, 10^−3^, and 10^−4^ were obtained following the conventional serial dilution method. From each dilution, aliquots were plated in triplicate on MRS agar plates.

The inoculated plates were incubated at 33 °C for 24 h, after which well-defined colonies were counted and expressed as colony-forming units per gram (CFU/g).

### Experimental design and statistical analysis

To determine statistically significant differences in CFU counts among treatments, a completely randomized design (CRD) was applied using a one-way analysis of variance (ANOVA). Statistical analyses were performed using Microsoft Excel 2016 on a Windows 12 operating system.

When a significant effect was detected in at least one treatment (p < 0.05), Fisher’s least significant difference (LSD) multiple range test was applied at a 95% confidence level to identify the statistical sources of parametric mean differences.

## RESULTS

### Growth curve of *Lactobacillus bulgaricus* in MRS broth

*Lactobacillus bulgaricus* was cultured under closed-system conditions in de Man, Rogosa and Sharpe (MRS) broth (HiMedia), where biomass growth corresponded to substrate consumption within the chemostat system.

The growth curve showed a progressive increase in biomass and turbidity over the 20 h incubation period. The onset of the exponential phase occurred at approximately 6 h, while the stationary phase was reached at around 18 h. Maximum growth was observed at 12 h, as shown in Figure 1. Consequently, this time point was selected for inoculating the black soldier fly larvae meal (BSFLM) and soybean meal mixtures for subsequent fermentation.

**Figure 1.**
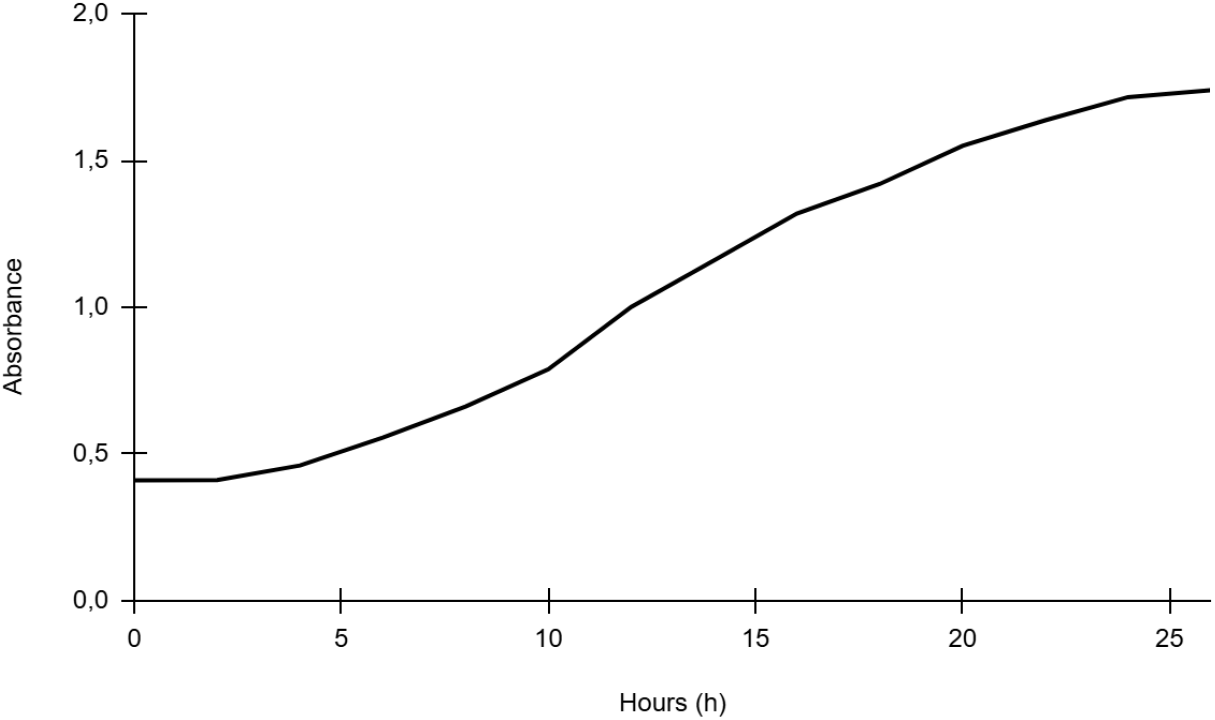
Growth curve of *Lactobacillus Bulgaricus* in MRS broth for 20 hours of incubation at 37 °C.

### Viable cell counting using the Neubauer chamber

Viable cells were quantified using the optimized Neubauer chamber method (Zhang et al., 2020). At the 1:10,000 dilution, the microscopic field shown in Figure 2 was obtained, representing the number of viable cells per 0.1 µL.

**Figure 2.**
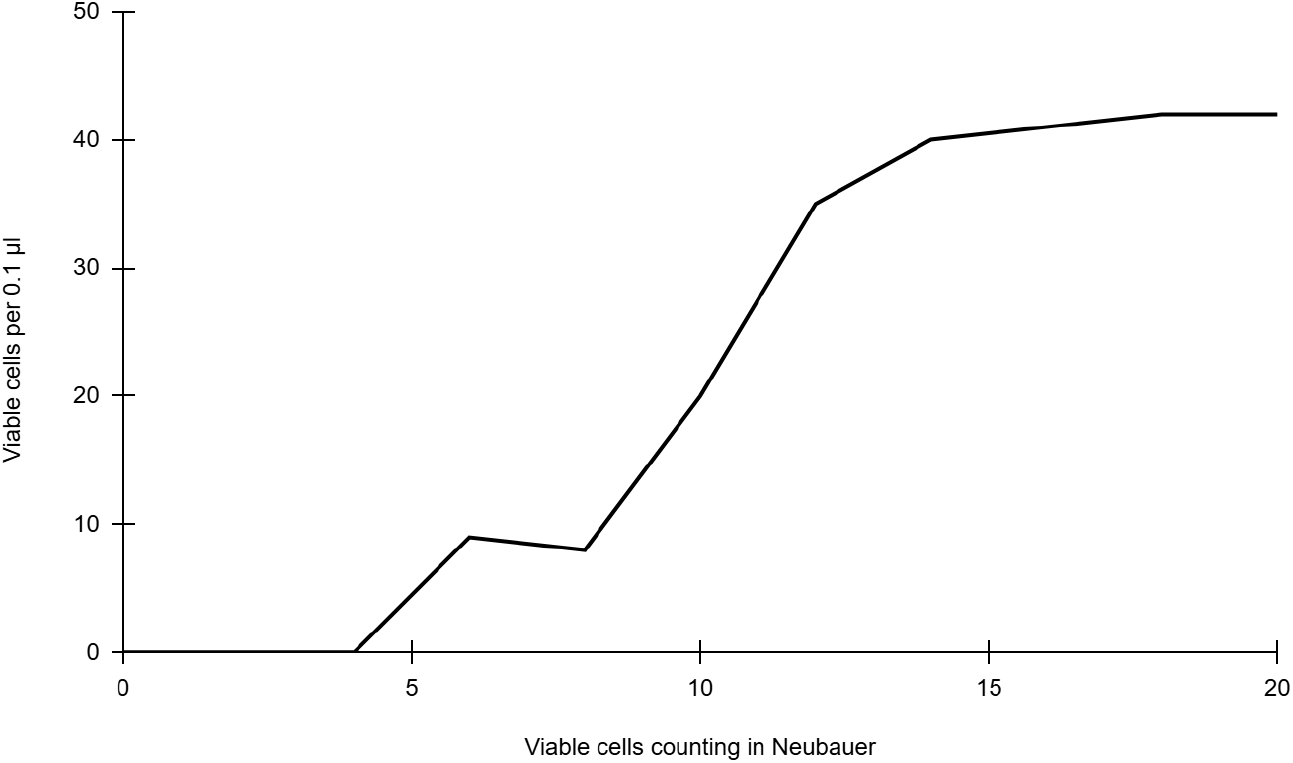
Viable cells per 0.1 µl from the dilution 1:10000.

Based on these results, it was calculated that the inoculum reached a concentration of 4.2 × 108 *Lactobacillus bulgaricus* cells per milliliter upon reaching the stationary phase. This value is consistent with previously reported growth data for *Lactobacillus* spp. under optimal culture conditions (Silva et al., 2018).

### Colony-forming unit (CFU) counts from serial dilution plate assays

Figure 3 shows a comparison of the mean CFU values obtained from triplicate plates for each formulation, highlighting the superiority of formulation 1 (pure BSFLM) over formulation 5 (pure soybean meal).

**Figura 3.**
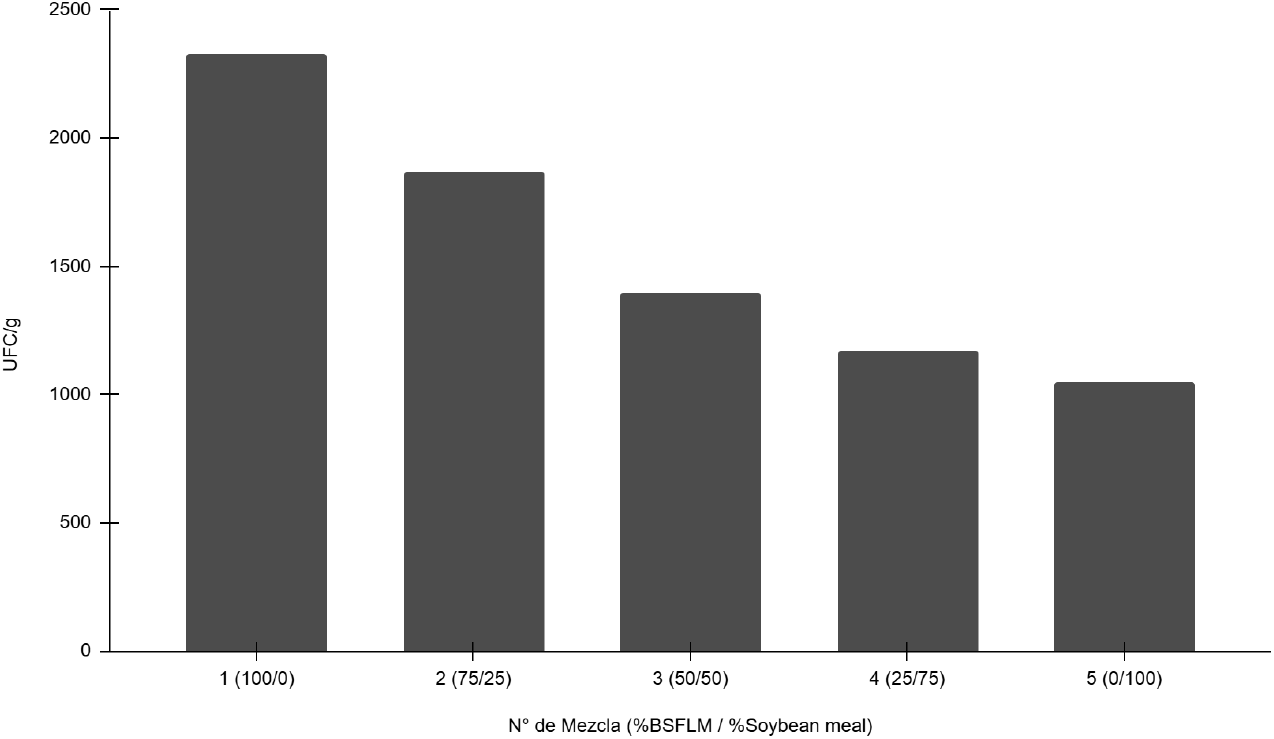
Media aritmética de las UFC/g de cada mezcla hecha.

As illustrated in Figure 5, formulations containing higher proportions of BSFLM consistently exhibited higher CFU counts compared to those based predominantly on soybean meal.

### Experimental design and statistical analysis

The experimental matrix is presented in Table 2, and the corresponding ANOVA results are shown in Table 3. The analysis yielded an F-statistic value of 130.14, exceeding the critical F value of 3.48, with a p-value of 1.40 × 10^−8^. These results clearly indicate the existence of statistically significant differences among at least one of the treatments, justifying the application of Fisher’s least significant difference (LSD) multiple comparison test.

**Table 2.**
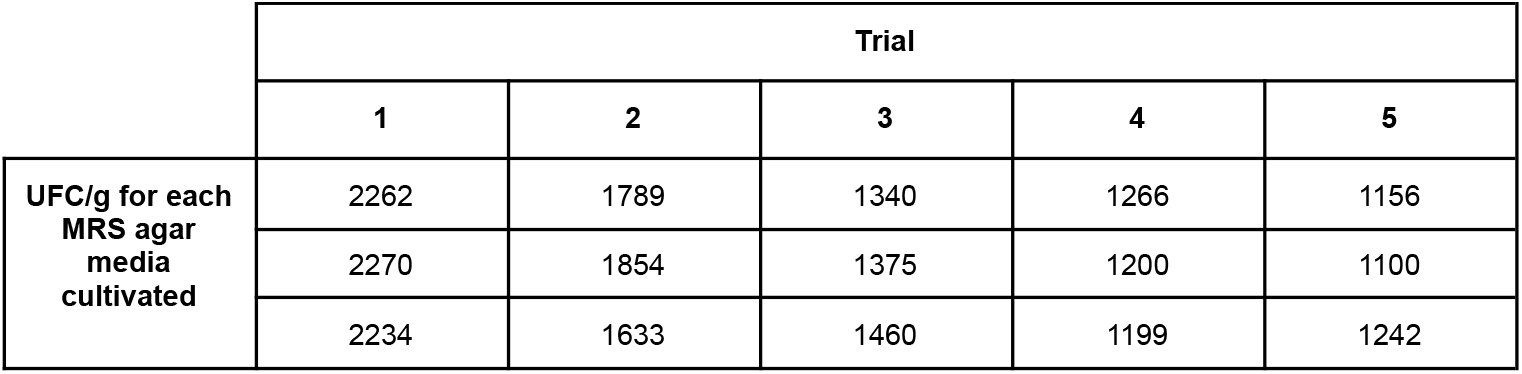
UFC/g for each trial tested.

**Table 3.**
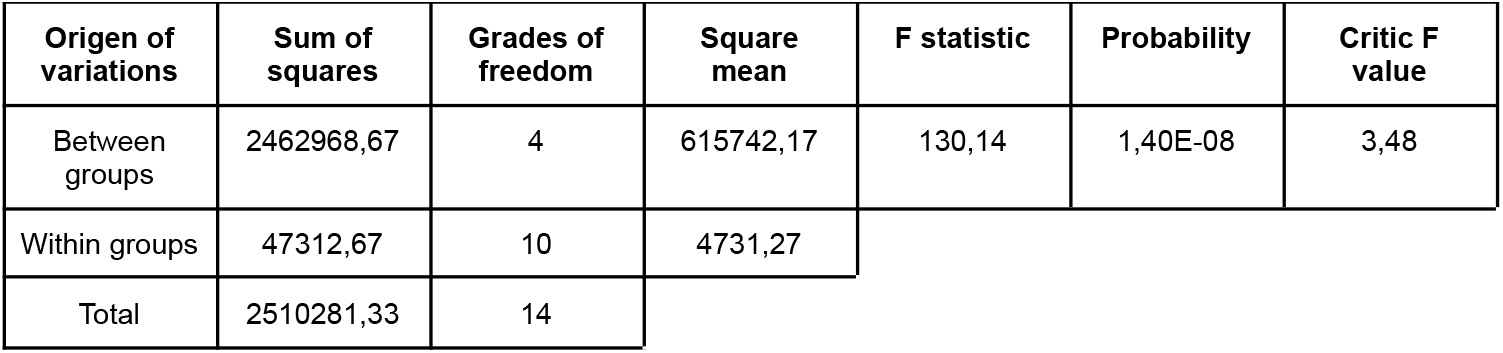
UFC/g in each trial done.

The Fisher-LSD test produced an LSD value of 125.14, which was compared against the absolute differences between treatment means, resulting in a total of 10 pairwise contrasts. Differences exceeding the LSD value were considered statistically significant. The analysis revealed that all treatments differed significantly from one another, except for treatments 4 and 5. These results indicate that BSFLM concentrations equal to or greater than 50% promote significantly higher *Lactobacillus bulgaricus* production, whereas concentrations equal to or below 25% do not significantly affect the maximum probiotic concentration.

Lactic acid bacteria (LAB) are probiotics that contribute to the nutritional value of foods and inhibit the growth of pathogens through the production of antimicrobial compounds, thereby benefiting gastrointestinal health in both humans and animals (Ansari et al., 2023). LAB, like other probiotics, can be used synergistically with insect-based feeds for terrestrial or aquatic animals intended for human consumption, conferring immunostimulatory properties that improve animal health (Anany et al., 2023; Hasan et al., 2023).

Previous studies have explored the use of black soldier fly larvae meal in combination with probiotics. For instance, Phaengphairee et al. (2023) evaluated this substrate combined with multi-probiotic formulations (*Bacillus subtilis, B. licheniformis*, and *Saccharomyces cerevisiae*), reporting increased nutrient digestibility, immunoglobulin A levels, and glutathione peroxidase activity, along with reduced pro-inflammatory cytokines in pigs.

During the fermentation process, no microbial growth other than LAB was observed, indicating clear dominance of *Lactobacillus* spp. in fermented BSFLM. Similar dominance was reported by Meng et al. (2023), who used the same substrate with a different *Lactobacillus* strain. This dominance is attributed to lactic acid production and the associated decrease in pH (3.99–6.60), which inhibits the growth of competing microorganisms, thereby extending product shelf life and potentially improving sensory characteristics. In the present study, the final pH of treatment 1 reached 5.20, whereas treatment 5 exhibited a pH of 6.20, which is consistent with the higher concentration of *Lactobacillus bulgaricus* observed at higher BSFLM proportions.

Despite achieving a considerable probiotic concentration in treatment 1, the viability of *Lactobacillus bulgaricus* decreased from approximately 8 log CFU/g in the initial inoculum to 3.35 log CFU/g in the fermented BSFLM after 48 h. This represents a reduction of approximately 4.65 log CFU/g, equivalent to a loss of ∼99.998% of viable cells. This finding is consistent with previous reports indicating that certain black soldier fly substrates, particularly prepupal or defatted meals, may not support the growth of specific *Lactobacillus* strains and may even exert inhibitory effects (Meng et al., 2023).

Several factors may account for this reduction, including the presence of natural antimicrobial compounds in the meal (e.g., chitin-derived peptides), limited availability of fermentable carbohydrates, and potential effects related to pH, water activity, residual lipid compounds, or insufficient fermentation time. Therefore, future studies should consider optimization designs incorporating variables such as *Lactobacillus* strain selection, water activity, fermentation time, and the developmental stage of the black soldier fly larvae used as substrate.

Consistent with these findings and current literature, the generation of feed formulations containing high concentrations of *Lactobacillus bulgaricus*, as achieved in the present study, represents a promising opportunity for innovation in sustainable livestock production systems (Chisoro et al., 2023; Biteau et al., 2024). Although in vitro or in vivo trials were not conducted to directly confirm positive health regulation effects, the present results provide a solid foundation supporting their potential.

Moreover, the substrate itself has been demonstrated to be a digestible and nutritionally adequate feed source for guinea pigs (*Cavia porcellus*), as reported by Reátegui et al. (2020), further reinforcing the biotechnological potential of BSFLM-based probiotic feeds within the proposed livestock production scenario.

## CONCLUSIONS

The present study demonstrated that fermentation mediated by *Lactobacillus bulgaricus* using pure black soldier fly larvae meal (BSFLM) as substrate resulted in a statistically significant higher concentration of this probiotic (p < 0.05) compared to the other formulations prepared with different proportions of soybean meal. The results provide evidence that BSFLM concentrations equal to or below 25% (w/w) do not lead to a significant increase in probiotic biomass.

On the other hand, although the pure BSFLM treatment exhibited the highest biomass concentration, an approximate reduction of 4.65 log CFU/g (equivalent to a loss of ∼99.998% of viable cells) was observed when comparing the initial MRS broth inoculum to the fermented product obtained after 48 h. This finding highlights the need to optimize the fermentation process prior to any potential scale-up. Furthermore, the results suggest that *Lactobacillus bulgaricus* may not be the most suitable lactic acid bacterium for BSFLM fermentation, or that the larval developmental stage is a critical factor to be considered in probiotic fermentations of this feed substrate. Nevertheless, the evidence provided in this study is expected to serve as a useful foundation for future research aimed at the production of probiotic feeds and the advancement of sustainable livestock systems.

## ACKNOWLEDGMENTS

The Catholic University of Santa María (Arequipa, Peru) provided the facilities and financial support necessary for the completion of this study. Also express special gratitude to Percy Alfredo Barbachán Medina, from the San Juan Bautista de La Salle educational institution (Arequipa, Peru), for his example and inspiration imparted to one of his students.

